# CRISPR-based diagnostics detects invasive insect pests

**DOI:** 10.1101/2023.05.16.541004

**Authors:** Pathour R. Shashank, Brandon M. Parker, Santosh R. Rananaware, David Plotkin, Christian Couch, Lilia G. Yang, Long T. Nguyen, N. R. Prasannakumar, W. Evan Braswell, Piyush K. Jain, Akito Y. Kawahara

## Abstract

Rapid identification of organisms is essential across many biological and medical disciplines, from understanding basic ecosystem processes and how organisms respond to environmental change, to disease diagnosis and detection of invasive pests. CRISPR-based diagnostics offers a novel and rapid alternative to other identification methods and can revolutionize our ability to detect organisms with high accuracy. Here we describe a CRISPR-based diagnostic developed with the universal cytochrome-oxidase 1 gene (CO1). The CO1 gene is the most sequenced gene among Animalia, and therefore our approach can be adopted to detect nearly any animal. We tested the approach on three difficult-to-identify moth species (*Keiferia lycopersicella, Phthorimaea absoluta*, and *Scrobipalpa atriplicella*) that are major invasive pests globally. We designed an assay that combines recombinase polymerase amplification (RPA) with CRISPR for signal generation. Our approach has a much higher sensitivity than other real time-PCR assays and achieved 100% accuracy for identification of all three species, with a detection limit of up to 120 fM for *P. absoluta* and 400 fM for the other two species. Our approach does not require a lab setting, reduces the risk of cross-contamination, and can be completed in less than one hour. This work serves as a proof of concept that has the potential to revolutionize animal detection and monitoring.

## Introduction

Accurate organismal identification is crucial for many biological and medical disciplines, from understanding basic ecosystem processes and determining how organisms respond to environmental change, to diagnosing diseases and detecting invasive pests (1, 2, 3). Unfortunately, traditional morphology-based identification approaches are often slow, require trained taxonomic specialists, and can result in unresolved or incorrect identifications when species are part of cryptic complexes (4, 5). Some of the first molecular approaches used Sanger sequencing (6) with Polymerase Chain Reaction (PCR) (7). Recently, rapid molecular identification methods have been developed and adopted as an important high capacity and accurate tool in efforts to identify species (e.g., 8, 9, 10, 11, 12). These approaches include multi-locus sequence typing (MLST) (13), real-time and droplet digital PCR (8,14), high-throughput sequencing (HTS) (15–18), loop-mediated isothermal amplification (LAMP) diagnosis (19), and SNPs iPLEX MassARRAY (20), and SHERLOCK (21). These methods have made it possible to detect target nucleic acids in diverse environmental DNA (eDNA) samples, allowing simultaneous detection of multiple target species, while decreasing the time needed to identify target taxa. However, protocols for these molecular approaches are still time consuming, expensive, or require a laboratory setting.

The recently developed CRISPR-Cas (clustered regularly-interspaced short palindromic repeats-associated system) (22) offers a novel and rapid alternative to other identification methods. CRISPR-Cas has been broadly applied to many disciplines in the life sciences, agriculture, biotechnology, and therapeutics. CRISPR-based approaches have also started to be implemented in diagnostics (21). They have the potential to combine the ease of use and cost efficiency of isothermal amplification with the ultrasensitivity of quantitative PCR but provide results in a much shorter amount of time (21, 23). CRISPR-Cas diagnostics combine isothermal preamplification of nucleic acid and programmable cleavage of nucleic acids, using guide RNA (gRNA) with Cas proteins, to attain attomolar sensitivity and single-nucleotide specificity (24, 25). The process takes advantage of the collateral cleavage (trans) activity from Class 2 Type V and Type VI Cas proteins, specifically Cas12a and Cas13a, to cleave a fluorescence resonance energy transfer (FRET)-based reporter resulting in fluorescence (21, 23). While CRISPR-Cas has been shown to be effective in the diagnosis of human and plant disease (26, 27), and in each case, unique genomic regions have been developed to serve as diagnostic targets. This approach requires detailed genomic information to identify candidate targets and population variation data to ensure consistency across populations, both of which can be expensive and labor intensive to acquire. Here we introduce an approach that can be applied to diverse taxa, including those lacking genomic resources, and in many cases using existing data.

The mitochondrial COI gene is one of the most frequently sequenced genes (28) due to its status as the primary ‘barcode’ sequence for animals. The BOLD database of COI sequences (29) currently includes over 12 million barcodes from over 250,000 species of animals from around the world. In order to maximize the utility of a CRISPR-based diagnostic approaches, it would be most effective if it is compatible with this widely sequenced gene. Here we present a CRISPR-Cas based diagnostic using the COI gene to provide highly accurate diagnoses. We demonstrate our approach on three moth species which are major agricultural pests globally: *Keiferia lycopersicella*, *Phthorimaea absoluta*, and *Scrobipalpa atriplicella*. We chose to work with these species because they are difficult to identify morphologically, because they are closely related (allowing for a test of sensitivity in similar target sequences), and because they are major pests that are collected as by-catch while using bulk traps in agricultural monitoring programs. We show that our CRISPR-based diagnostic platform can be utilised as a generalised molecular diagnostic method for any animal and has the potential to revolutionize organismal identification techniques.

## Results

### RPA-CRISPR-Cas12a assay design and optimization for insect diagnosis

We designed our assay to detect three different closely related moth species that are morphologically difficult to distinguish (Fig. 1a) using recombinase polymerase amplification (RPA) of COI before Cas12a detection. First, we designed and optimized a two-pot assay in which RPA and CRISPR-Cas12a reactions were conducted separately (Fig. 1c). The specificity, sensitivity (LoD), endpoint detection (fluorescence and lateral flow strip) and field sample testing for the two-pot assay worked for all three species. We successfully identified all three species using the two-pot assay with 100% accuracy, with detection limits of up to 120 fM for *P. absoluta* and 400 fM for *K. lycopersicella* and *S. atriplicella*. Later, a one-pot assay was developed by combining RPA and CRISPR-Cas reactions in a single tube (Fig. 1d). We report that the RPA-CRISPR-Cas12a one-pot assay on extracted DNA achieved 100% accuracy for *P. absoluta* and *K. lycopersicella*, but it achieved 94.11% accuracy for *S. atriplicella* with a 5.89% false negative rate. The sensitivity and diagnostic specificity (field samples) of the one-pot assay were determined for target species samples.

**Figure 1.**
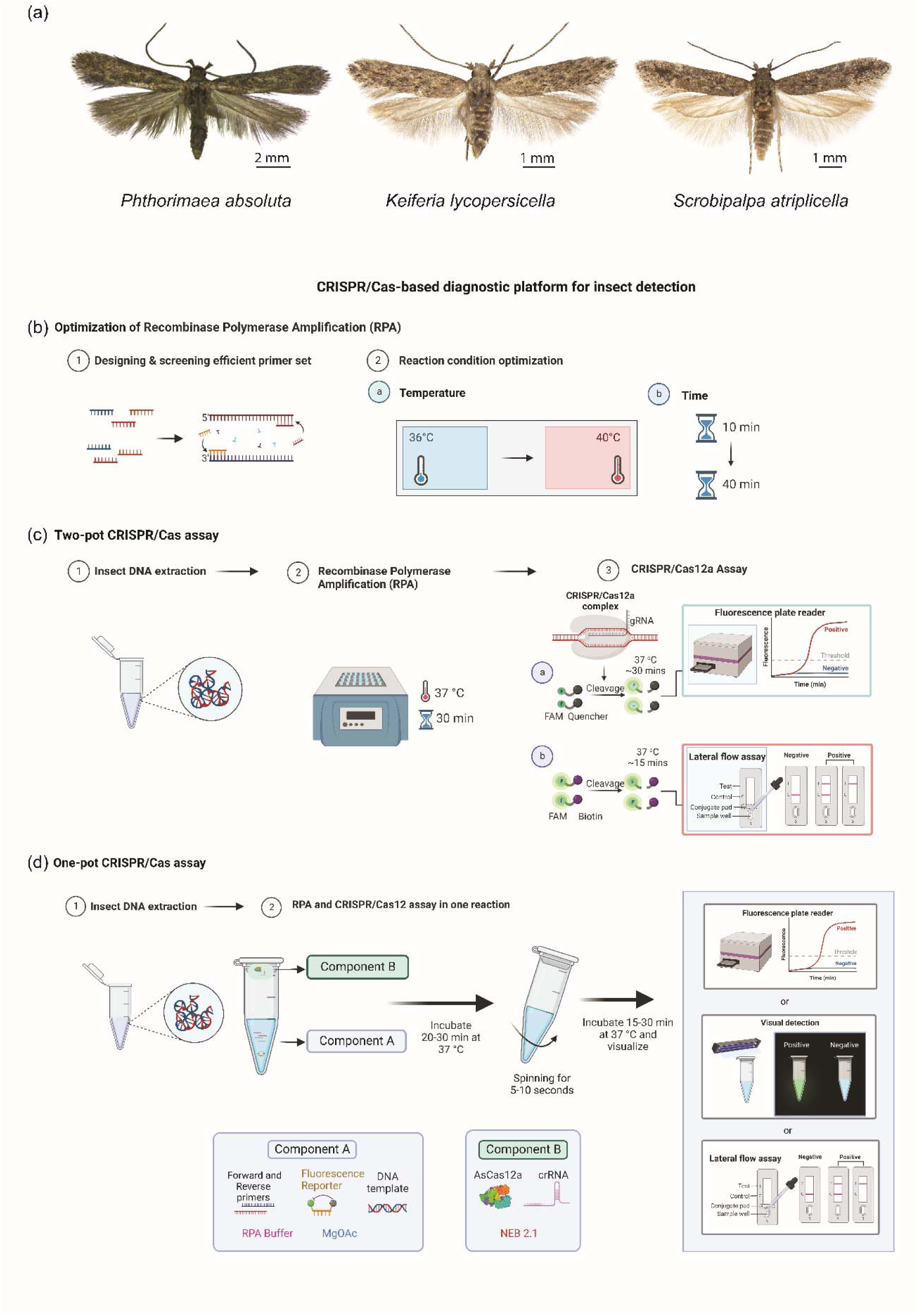
The CRISPR/Cas-based diagnostic platform for insect detection. (**a**) Target species included in the present study as models for developing the CRISPR/Cas assay. (**b**) Recombinase polymerase amplification (RPA) optimization scheme. (**c**) The two-pot CRISPR/Cas assay. (**d**) The one-pot CRISPR/Cas assay. DNA is extracted from samples with a column-based kit method (DNeasy Blood & Tissue Kit). (Fig 1b, c, d were “Created with <u>BioRender.com</u>”).

### RPA primer design, screening and optimization

Instead of designing species-specific primers for each species, we aimed to design a single set of RPA primers that will amplify the target region for all three target species (Fig. 1b). We constructed four forward primers (F1–4) and four reverse primers (R1–4) using TwistDx manufacturer guidelines (*SI appendix,* Table S1). Primers were 30–35 nucleotides long and the amplicon target length was 200–300 base pairs. A total of 16 combinations of primers (F1:R1-4, F2:R1-4, F3:R1-4, F4:R1-4) were screened and the RPA-F4 and RPA-R4 primer pair was selected because it successfully amplified all three species (*SI Appendix,* Figs. S1–S3). When RPA was performed at four different time intervals (10, 20, 30, 40 mins), we found that all temperatures showed amplification success. However, the trans-cleavage activity of Cas12a is at 37°C therefore we selected 37°C as the optimum temperature for further experiments (*SI Appendix,* Fig. S4). For RPA, amplification was satisfactory at 20, 30, and 40-minute intervals. We therefore chose the median value, 30 mins, as a sufficient time for further experiments (*SI Appendix,* Fig. S5).

### Cas12a enzyme, crRNA selection and specificity testing

In the present study, we designed two gRNAs for each insect species based upon the TTTV Protospacer Adjacent Motif (PAM) region within the designated RPA amplicon region (*SI Appendix*, Fig. S6). Subsequently, we tested the nucleic acid detection properties of three different Cas12 enzymes, *Lachnospiraceae bacterium ND2006* Cas12a (LbCas12a), *Acidaminococcus* sp. Cas12a (AsCas12a) and *Eubacterium rectale* Cas12a (ErCas12a), for evaluating the reporter cleavage activity with two gRNA spacer sequences for their corresponding activator molecules (target DNA sequences) (Fig. 2a–d). We observed that in *P. absoluta* and *S. atriplicella*, AsCas12a detection is substantially faster than that of ErCas12a and LbCas12a. Furthermore, AsCas12a exhibited a more than 20-fold increase in signal within 20 min (Fig. 2a, c). For *K. lycopersicella*, both AsCas12a and ErCas12a showed higher fluorescence at 20 min (Fig. 2b). We therefore selected AsCas12a for downstream analyses, as it worked best for these particular guides.

**Figure 2.**
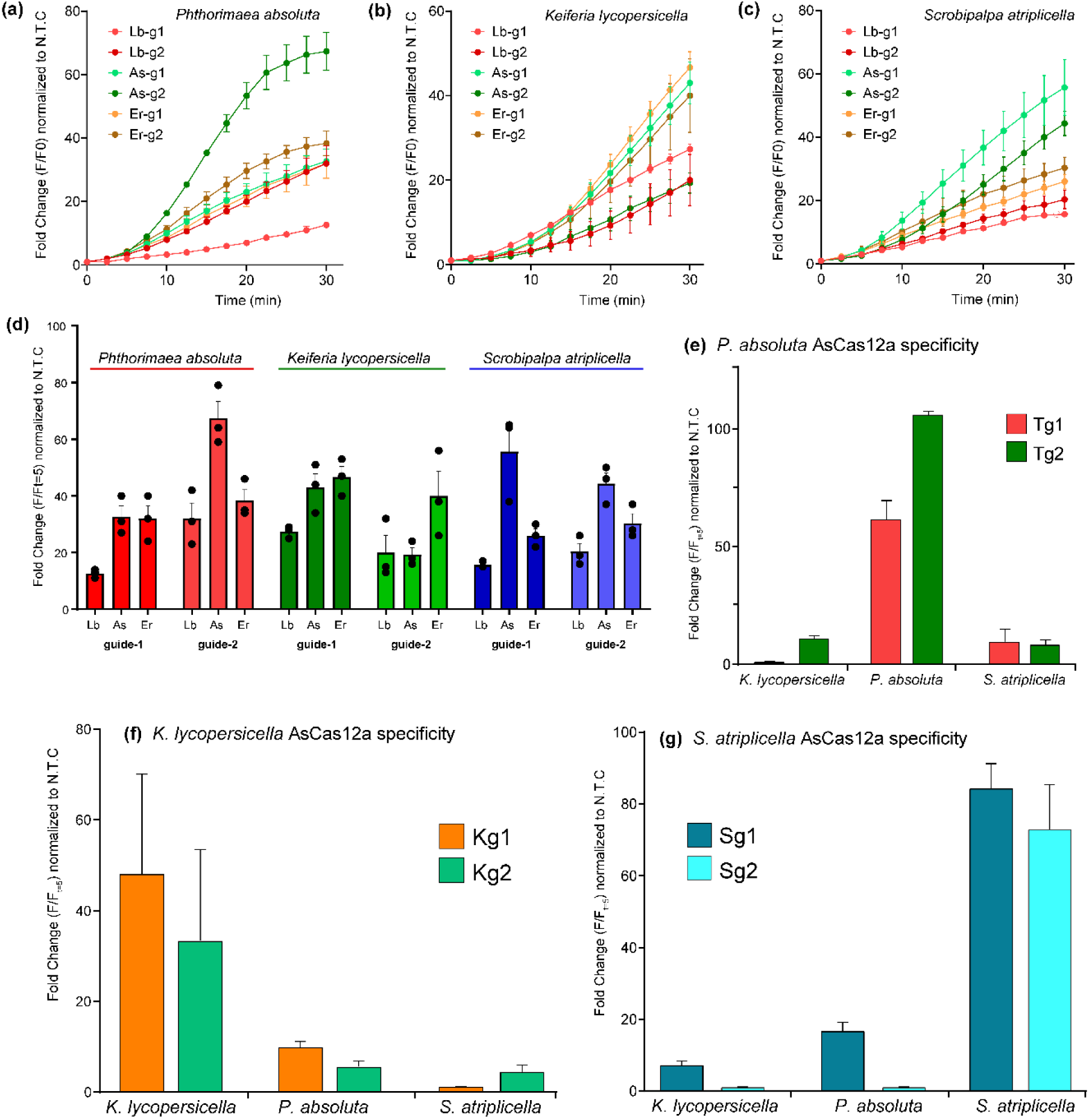
Screening and analytical specificity testing of Cas12a orthologs and guide RNA for the detection of insect species (a-c). Graphs depicting the fold change in fluorescence intensity with respect to the no-template control (NTC) against time for the detection of a synthetic DNA resembling the COI gene in *P. absoluta* (Pa)*, K. lycopersicella* (Kl), and *S. atriplicella* (Sa) with Lb, As, and Er Cas12a orthologs and two different guide RNAs for each ortholog. (d) Bar graphs showing fold change with respect to the NTC for the data in panels (a-c) at time t=30 min. (e-g) Specificity testing of the CRISPR-Cas12a assay against highly similar moth species using two gRNAs designed for each species.

To determine the specificity of our assay, we compared the species-specific gRNAs designed for *P. absoluta*, *K. lycopersicella* and *S. atriplicella*. We considered only these three species because they are difficult to identify and also occur as by-catch while monitoring using bulk traps. Limited cross-reactivity was observed for *P. absoluta* (Fig. 2e), *K. lycopersicella* (Fig. 2f), and *S. atriplicella* (Fig. 2g) for their particular gRNAs. However, there was a difference in fluorescence among the gRNAs, with three showing relatively high specificity: Tuta-crCOI-2 (Tg2) for *P. absoluta*, KP-crCOI-1 (Kg1) for *K. lycopersicella* and gRNA SP-crCOI-2 (Sg2) for *S. atriplicella* (Fig. 2e– g; *SI Appendix,* Table S2).

### Analytical sensitivity of two-pot RPA-CRISPR-Cas12a assay

The limit of detection (LoD) for the RPA-CRISPR-Cas12a assay was determined by using the PCR product of the target DNA template to make a serial dilution (ranging from 10 nM to 1 aM) followed by an isothermal pre-amplification RPA step. Results of the RPA amplification analysis by agarose gel electrophoresis showed that in *P. absoluta, K. lycopersicella* and *S. atriplicella* the concentration of 10 fM detected positive results, while after this concentration, bands were fainter (*SI Appendix,* Fig. S7). However, these results were not consistent with the RPA-CRISPR-Cas12a. Therefore, first an estimated LoD was determined based on the lowest concentration having 3/3 of replicates showing at least a three-fold increase in fluorescence signal compared to that of the no-template control (NTC) in 30 min (Fig. 3a–c). For *P. absoluta*, a concentration of 10 fM exhibited the lowest limit of positive detection with 3/3 replicates. For *K. lycopersicella* and *S. atriplicella*, this limit was at 100 fM. To further confirm the exact LoD, we selected concentrations around the lowest limit of detection and tested five replicates. For *P. absoluta,* we tested 120 fM, 80 fM, 60 fM and 40 fM and found the LoD to be 120 fM within 30 min for a CRISPR reaction. Similarly, we tested four different concentrations for *K. lycopersicella* and *S. atriplicella* (50 fM, 100 fM, 200 fM, 400 fM) and estimated the LoD to be 400 fM. All five replicates showed signal of positive fluorescence (Fig. 3d–f) and were visible to the naked eye at the indicated LoD with blue light (Fig. 3g–i).

**Figure 3.**
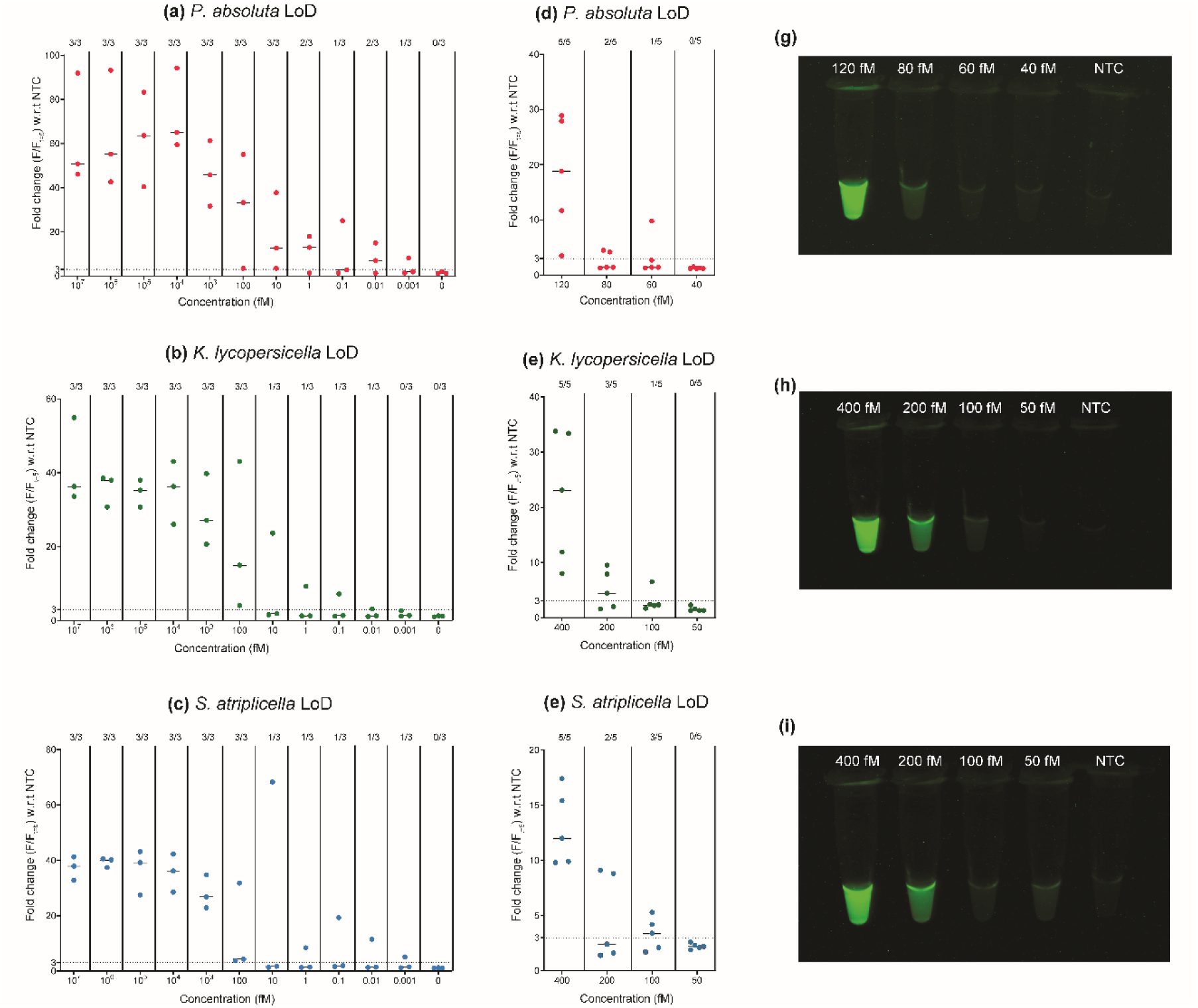
Limit of detection (LoD) tests for the CRISPR-Cas12a assay. (a-f) Fluorescence intensities in fold change with respect to the no-template control (NTC) in serially diluted samples containing PCR-purified COI gene fragments against the best gRNAs identified for *P. absoluta* (Tuta-crCOI-2)*, K. lycopersicella* (KP-crCOI-1), and *S. atriplicella* (SP-crCOI-2). **d-f** (n=3) two out of three of samples detected in **a-c** (n=5) were subjected to serial dilutions around the estimated LoD for the COI gene. (g-i) Images taken under blue light of the samples detected in d-f, respectively.

After testing specificity and sensitivity for analytical samples, we sought to evaluate the RPA-CRISPR-Cas12 assay with diagnostic DNA extracted from field samples of all three species. Samples were randomly selected and the person performing extractions was not told which samples corresponded to which species. Three experiments were performed, each consisting of 17 individuals of one of the target species, 6 randomly chosen members of the other two non-target species, and an NTC (Fig. 4a–c) and the results were interpreted using a fluorescence-based reporter and lateral flow-based detection. The criterion for a positive sample in fluorescence-based reporter detection is a change in fluorescence signal of 3-fold or greater compared to the NTC within 30 min. Out of a total of 51 specimens tested in separate diagnostic specificity assays (Seventeen each of *P. absoluta* (Fig. 4a), *K. lycopersicella* (Fig. 4b), and *S. atriplicella* (Fig. 4c)), the RPA-CRISPR-Cas12a assay detected DNA in all 51 samples, whereas all 21 negative and control samples showed no signal, indicating 100% accuracy. We also performed a lateral flow-based, 15-minute detection assay for the same samples, and its results showed 100% agreement with the fluorescence-based reporter assay (Fig. 4g– i).

**Figure 4.**
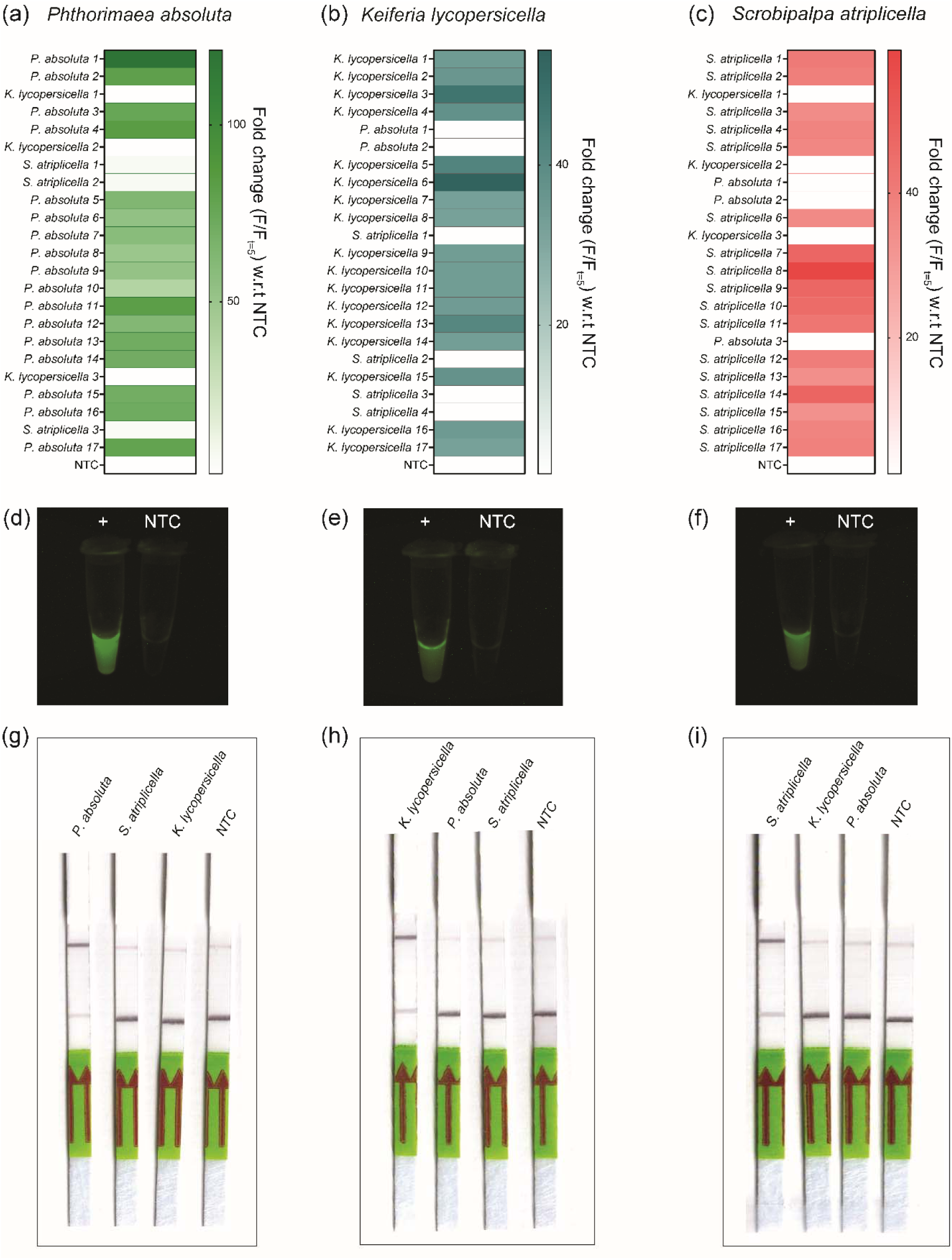
Specificity of the two-pot CRISPR assay. (a-c) Fluorescence intensity in fold change with respect to the no-template control (NTC) for detecting genomic DNA. (a) Detection of *P*. *absoluta* using gRNA Tuta-crCOI-2. (b) Detection of *K. lycopersicella* using gRNA KP-crCOI-1. (c) Detection of *S. atriplicella* using gRNA SP-crCOI-2. (d-f) Representative images taken under blue light of positive samples detected in a-c, respectively, alongside corresponding NTCs. (g-i) Representation of lateral flow assay testing of the three species. Full details are in *SI Appendix* Fig. S10.

### Development of a one-pot RPA-CRISPR-Cas12a detection method

We tested trans-cleavage activity by AsCas12a in a one-pot assay by combining both RPA and CRISPR-Cas12a reagents simultaneously using serial dilutions of *P. absoluta* genomic DNA. The AsCas12a enzyme showed high sensitivity and began trans-cleavage before preamplification of the target region (*SI Appendix,* Fig. S11) and previous studies showed that many CRISPR reagents inhibit RPA reaction when added together (Wang et al. 2019). To address this issue, we adopted and developed a method that combines the approaches of Ding et al. (2020), Sun et al. (2021) and Xiong et al. (2022). Specifically, Cas12a with its gRNA and NEB2.1 buffer were loaded separately in a tube lid before being mixed with the RPA reaction. After spinning, we observed that approximately 5 µL of the CRISPR/Cas12a reaction mixture on the lid of the tube was small enough to be held on the lid by surface tension. We tested 10-, 20- and 30-minute incubation times at 37°C before spinning for three test species. Results showed different fluorescence signals for different species (*SI Appendix,* Fig. S12). Thirty minutes was found to be the optimal incubation time, with a more than 20-fold increase in signal for all species, and therefore we used this spin time for further experiments to minimize the chance of false negatives.

Further, we determined the limit of detection for the one-pot assay using a ten-fold serial dilution for *P. absoluta*, *K. lycopersicella*, and *S. atriplicella* (Fig. 5a–c, respectively). The one-pot assay confirmed that we can detect genomic DNA of *P. absoluta* in concentrations as low as 10^−7^ ng/μl, and *K. lycopersicella* and *S. atriplicella* in concentrations as low as 10^−5^ ng/μl. We evaluated diagnostic specificity of the one-pot assay with the same insect samples used to test diagnostic specificity in the two-pot assay. In the one-pot assay, we also were able to correctly detect all 17 samples of *P. absoluta* and 17 samples of *K. lycopersicella* (Fig. 5e) without false positives or negatives (*SI Appendix,* Figs. S13, S14), and we were able to do this for *S. atriplicella* in 16/17 samples (Fig. 5f, *SI Appendix,* Fig. S15). The entire one-pot assay requires only 50–60 mins while using a fluorescence-based reporter. On the basis of the sample dataset evaluated for diagnostic specificity, we report the RPA-CRISPR-Cas12a one-pot assay on extracted DNA as having 100% accuracy with false positive and false negative rates of 0% for *P. absoluta* and *K. lycopersicella*, and 94.11% accuracy with a 5.89% false negative rate for *S. atriplicella*.

**Figure 5.**
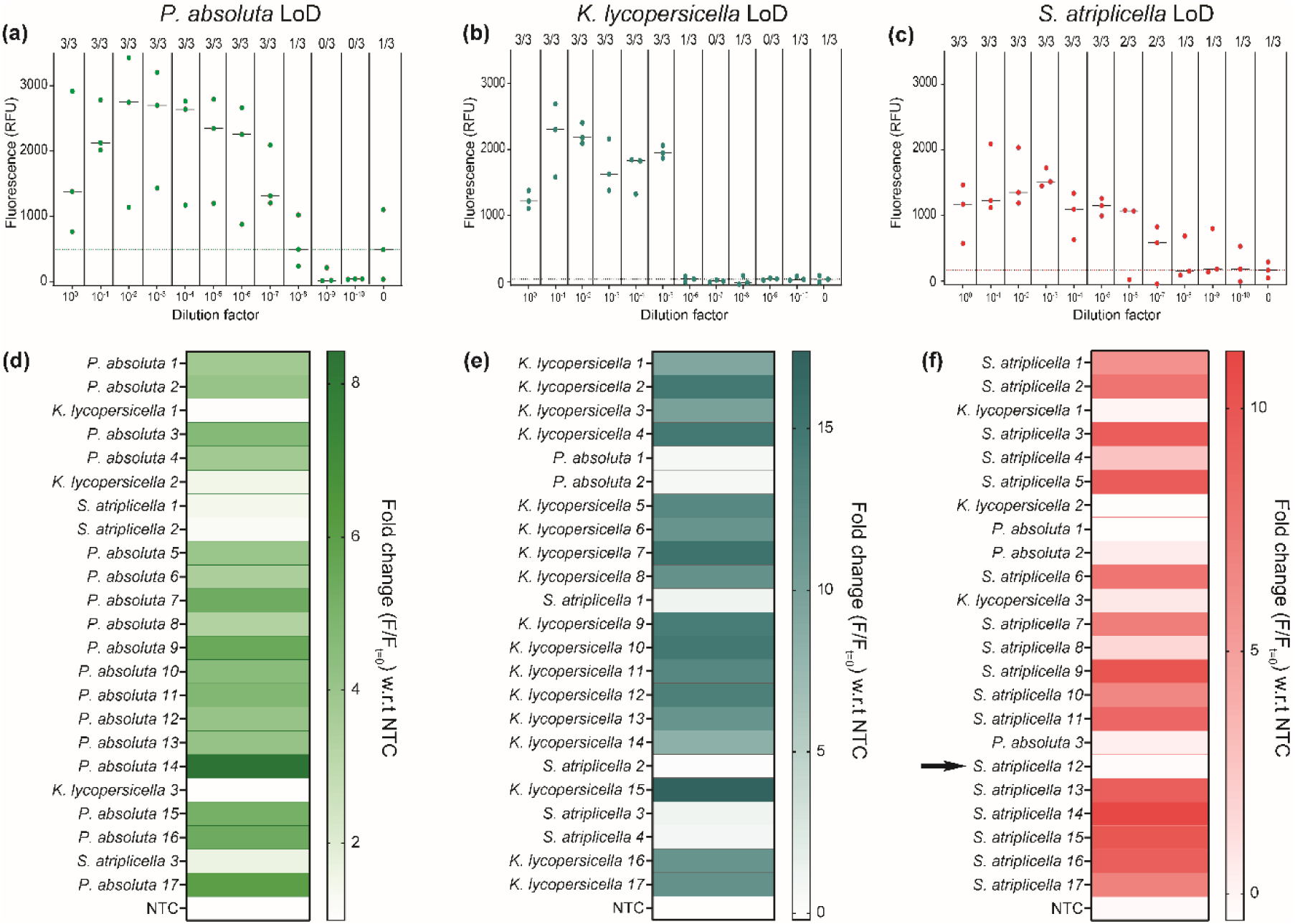
Testing the specificity of the one-pot CRISPR detection assay. (a-c) Fluorescence intensities in RFU in 11 different serially diluted samples containing gnomic DNA against the best gRNA’s identified for *P. absoluta* (Tuta-crCOI-2)*, K. lycopersicella* (KP-crCOI-1), and *S. atriplicella* (SP-crCOI-2) (n=3). (d-f) Fluorescence intensity in fold change with respect to NTC for detecting genomic DNA. (d) Detection of *P. absoluta*. (e) Detection of *K. lycopersicella.* (f) Detection of *S. atriplicella*. The arrow in Fig. 5f indicates a false negative.

## Discussion

Rapid detection of organisms is critical for many disciplines, including ecological monitoring, detecting bacterial and viral spread, and preventing the introduction of invasive pests. Advances in molecular biology have led to many new approaches, including ddPCR, qPCR, TaqMan real-time PCR, and high-throughput sequencing (18, 34, 35). None of these methods have taken over as the standalone diagnostic because they each have their own advantages and disadvantages. Still, new molecular diagnostic methods continue to be developed. Recently, nucleic acid detection using CRISPR-based detection platforms such as DETECTR (23), SHERLOCK (21), and ENHANCE (36) have emerged as alternatives to traditional RT-qPCR assays because of their greater sensitivity, specificity, and speed (24, 27, 37). However, these technologies have been utilized primarily for disease diagnosis (38, 39, 40, 41), and very few have been applied to the detection of non-model organisms. Although there are more than 1.5 million described animal species on Earth (42), CRISPR technology has not been developed widely for the detection of non-model animal taxa.

We designed species-specific gRNAs using the COI gene, a marker that has the most available sequence data among animal species (29). COI is also an ideal target locus because it is an abundant and robust mitochondrial gene (28). When we tested the specificity of our RPA-CRISPR-Cas12 assay on our closely related test species, all were detected with 100% accuracy and showed no false positives. Our RPA-CRISPR-Cas12a method can detect cryptic species in less than an hour after DNA extraction, results which are congruent with studies that utilize CRISPR for the detection of malaria (40) and COVID-19 (36). Although implementation of our approach requires having a COI sequence of the target taxon, our assay is easily applicable to any taxon because of the availability of reference COI data for many species in databases such as BOLD (29), making it easy to obtain the target sequence if a reference is unavailable. While the COI gene sequence is relatively invariable across populations of the same species (43), there are cases where this is not true (e.g., 44). Given the specificity of our RPA-CRISPR assay, we recommend sequencing COI for the target population of the target taxon whenever possible.

We first designed, screened, and tested different sets of common RPA primers and identified RPA-F4 and RPA-R4 as the best primers to amplify selected species under isothermal assay conditions (37°C). Preamplification methods like RPA have stringent reaction condition requirements and it is important to select the isothermal amplification reaction compatible with Cas12a. While LbCas12a is a frequently used variant for CRISPR diagnosis in plants (45) because of its high efficiency at 37°C (46), there has not been a single study thus far that outlines the best Cas12a variant for insect diagnosis. Therefore, we tested three different variants (AsCas12a, ErCas12a, LbCas12a) and incorporated AsCas12a into our protocol because it performed better than the latter two. Reasons remain unknown as to why the latter two variants performed poorly, but we predict that it may be due to specific properties such as temperature stability or enzyme kinetics of the variant (47).

Our RPA-CRIPSR approach is a one-pot assay where RPA and CRISPR reactions occur in the same tube. We developed this protocol from a two-pot assay that involved additional steps. A one-pot isothermal amplification coupled with CRISPR/Cas systems are the preferred choice for next generation molecular diagnostics because it reduces the risk of contamination and speeds up processing time (47, 48). Our assays showed very high sensitivity (10^−7^ ng/μl for one-pot and 120 fM for two-pot) compared to a recently developed RT-PCR assays, where LoD is typically ≥ 0.01 ng/µl (8) and droplet digital PCR assays with LoDs of 0.001ng/µL (49) (Data only for *P. absoluta* as comparative data are unavailable for the two other species). Our assay is rapid; the RPA reaction takes 30 minutes of incubation and 15 minutes to obtain lateral flow results. The entire process can be completed in less than one hour.

There are several areas where our assay can be improved. First, although we identified AsCas12a as the best variant for our approach, our results suggest that with further optimization, ErCas12a can potentially also be developed as a novel Cas12a-based detection platform. We therefore recommend future studies to examine whether ErCas12a can be used as an alternative to AsCas12a. Another area of potential improvement is in the DNA extraction step. We used a Qiagen DNeasy extraction kit, but there are many other approaches such as OmniPrep™ Genomic DNA isolation kit, Sigma’s GenElute™-E Single Spin Tissue DNA Kit etc. but there are no studies that compare these to Qiagen and prove that they’re faster. Future studies should examine how extraction protocols can improve and speed up our molecular diagnostic. Thirdly, our approach was designed in a lab. Although this can easily be replicated in the field, creating a portable “field kit” with all reactions and tubes would be ideal for immediate deployment and efficiency of identifying target taxa. Finally, developing a lyophilized CRISPR version (see 22), would improve the assay as it not only reduces reaction time further but also simplifies the need to set up a CRISPR reaction.

## Materials and Methods

### Insects

We selected three morphologically similar moth species in the same family (Gelechiidae), that are difficult to identify (Fig 1a), in order to test the efficacy of our CRISPR approach. The three species are: (a) Tomato leaf miner moth, *Phthorimaea absoluta*, a highly invasive moth pest that damages tomato, eggplant, pepino, pepper, and many other crops. The larva feeds voraciously on tomato plants, producing wide galleries in leaves, burrowing in stems and destroying fruit. Crop yield loss can reach 100% (50, 51). The moth has invaded South America, and is now present in the Caribbean islands and is a major threat to US agriculture (52). (b) Tomato pinworm*, Keiferia lycopersicella*, an important pest of tomatoes in North and Central America often confused with other microlepidopteran pests. The feeding damage is almost identical to that of *P. absoluta* and difficult to diagnose in the field (53). (c) Goosefoot groundling moth, *Scrobipalpa atriplicella*, another invasive species introduced to North America from Eurasia and recently reported to be an important pest on quinoa in Canada (54). The larva of *S. atriplicella* feeds on foliage, bores into stems, and also feeds directly on the panicle of seeds, causing significant damage to quinoa (54).

### DNA extraction, PCR and sequencing of insect samples

We received field-captured and laboratory-raised adults and larvae for all three moth species, as well as pupae of *P. absoluta* (*SI Appendix,* Table S6). Upon receiving specimens, we stored them at –80°C for at least two weeks before working with them, in order to kill any parasites or parasitoids potentially inside the specimens.

Genomic DNA was extracted from the whole body of specimens using a Qiagen DNeasy Blood and Tissue Kit (Qiagen, Valencia, CA, USA), following the manufacturer’s protocol with slight modifications to increase DNA yield from insects. Specimens were first homogenized by adding 180 µL ATL buffer, 20 µL proteinase K, and two 2.4 mm metal beads (VWR 500g x 2.4mm Metal). The microcentrifuge tubes were placed in a Mini-BeadBeater-96 homogenizer (Biospec) for 90 seconds at 1900 rpm. Samples were then incubated overnight at 56°C to lyse the cells. To elute the DNA, we added 50 µL of AE Buffer directly to the spin column filter and allowed it to incubate for 15 minutes at room temperature. This was repeated for a total volume of 100 µL of eluent. Extracted DNA was quantified using a Qubit 4.0 (Invitrogen, Thermo Fisher Scientific) broad-range double-stranded DNA assay. Isolated DNA was stored at 4°C for short term storage and at –20°C for long term storage.

The identification of the specimens was confirmed by sequencing the mitochondrial cytochrome oxidase I gene (COI). Extracted DNA was amplified using primers targeting COI (Invitrogen, LCO (forward primer): GGTCAAATCATAAAGATATTGG; HCO (reverse primer): TAAACTTCAGGGTGACCAAAAAATCA) via a 20 µL PCR that utilized the following: 10 µL Onetaq Quick-Load 2x master mix (New England Biolabs), 10 µg bovine serum albumin, 0.2 µM of each primer, 2 µL DNA, and 6.7 µL of PCR-grade water. Amplified DNA was shipped to Eurofins Genomics for Sanger sequencing. The resulting DNA sequences were checked for similarity using NCBI BLAST and trimmed using Geneious software (Geneious 11, Geneious Prime).

### Recombinase polymerase amplification (RPA) primer design

We elected to use RPA because it requires only two primers, has a low incubation temperature (35–42°C) and is more effective for point-of-care diagnosis compared to other isothermal amplification techniques (55) (Fig.1b). The 650 bp portions of COI we obtained from each of the three target moth species, as described in the previous paragraph (See also *SI Appendix,* Table S5), were used to create a consensus sequence. After polymorphic sites between species were identified, we designed RPA primers following the TwistAmp® assay design manual guidelines (http://www.twistdx.co.uk) using SnapGene software (GSL Biotech; available at www.snapgene.com); primers are listed in *SI Appendix*, Table S1. Following these protocols, we selected primer lengths to be 30–35 bp, amplicon length to be 100–500 nucleotides, and GC content to be 30%–70%. We also used BLAST to confirm that there were no significant similarities to COI sequences of other insect species. Our primers were synthesized by Integrated DNA Technologies (IDT).

### Recombinase polymerase amplification (RPA) and primer screening

The RPA assay was conducted to compare the performance of the four primer sets in amplifying the COI gene from all three species using manual guidelines (http://www.twistdx.co.uk). We conducted experiments in volumes of 50 µL using a TwistAmp® Basic kit. In each reaction, 29.5 µL of rehydration buffer, 2.4 µL (10 µM) of each forward and reverse primer, 3 µL of DNA, 10.2 µL of nuclease-free PCR-grade water and 2.5 µL (280 mM) of magnesium acetate were added to enzyme pellets, followed by incubation at 37°C for 20 min in a thermocycler (Bio-rad CFX96 Real-Time system). In all the assays, the DNA and magnesium acetate were added before incubation. All reactions were prepared in a laminar flow hood to reduce the chance of carry-over DNA contamination from previous reactions due to aerosolization.

We selected the best primer set which can successfully amplify the target region of all three species and later checked the optimal conditions for the RPA reaction. The samples were incubated at different reaction temperatures (36, 37, 38, 39, 40°C) and for different durations (10, 20, 30, 40 min). Reactions were halted by placing samples at or below 4°C. Amplified products were visualized by electrophoresis on a 2% v/w agarose gel stained with SYBR Green.

### Protein expression and purification

Bacterial expression plasmids for AsCas12a, and LbCas12a (Addgene #90095 and #90096, respectively) were obtained from the Zhang Lab (Broad Institute). The ErCas12a (Addgene #174685) expression vector was obtained from the Jain Lab at the University of Florida, Gainesville, FL, USA. The expression and purification procedures for these three nucleases were performed following the protocol of Nguyen et al. (48) with modifications. In brief, plasmids were transformed into Rosetta2(DE3) Plyss Singles^TM^ competent cells (Millipore Sigma, Catalog #70236), and individual colonies were chosen and optimized for protein expression during the following day. Selected colonies were inoculated in 2 liters of Terrific Broth (ThermoFisher, Catalog #H26824-36) at 37°C until OD = 0.6 – 0.8, followed by the addition of 500 mM IPTG (β-D-1-thiogalactopyranoside, Gold Biotechnology, Catalog #I2481C25) for protein expression and incubation at 18^°^C for 15-18 hrs. Next, *E. coli* cells were harvested, suspended in lysis buffer (500 mM NaCl, 50 mM Tris-HCl, PH = 7.5, 0.5 mM TCEP, 20 mM imidazole, 1 mM PMSF, 0.25 mg/mL lysozyme, 5% glycerol), and disrupted by sonication for 30 minutes. The cell lysate was centrifuged at 40,000 x g for 30 minutes. The supernatant was filtered through a 0.22 µm filter (Millipore Sigma, Catalog #SLGP033RS) before being injected into a Fast Protein Liquid Chromatography (FPLC) system (Bio-rad) that was connected to a HisTrap FF 5 mL column (Cytiva, Catalog #17525501) pre-equilibrated with a wash buffer (500 mM NaCl, 50 mM Tris-HCl, PH = 7.5, 0.5 mM TCEP, 20 mM imidazole). After another wash step was used to eliminate contamination, the column was eluted using an elution buffer (500 mM NaCl, 50 mM Tris-HCl, PH = 7.5, 0.5 mM TCEP, 300 mM imidazole, 5% glycerol). The elute was pooled and subjected to TEV protease treatment overnight at 4^°^C to remove the Maltose Binding Protein tag (MBP). The protein was then equilibrated with buffer A (150 mM NaCl, 20 mM HEPES, PH = 7, 0.5 mM TCEP, 5% glycerol) at 1:1 ratio prior to being injected into a HiTrap Heparin HP 1 mL column (Cytiva, Catalog #17040601) followed by gradient elution from buffer A to buffer B (2000 mM NaCl, 20 mM HEPES, PH = 7, 0.5 mM TCEP, 5% glycerol). Eluted fractions were analyzed using SDS-PAGE gels, and the purest fractions were collected, concentrated, exchanged into a final buffer (500 mM NaCl, 50 mM Tris-HCl, PH = 7.5, 0.5 mM TCEP, 5% glycerol), flash frozen, and stored at -80^°^C.

### gRNA and Cas12a nucleases selection

The guide RNAs (gRNAs) for Cas12a consist of two regions: a direct repeat (DR) region which helps in tethering the crRNA to the Cas protein, and a spacer region which is directly involved in target DNA recognition. We designed different gRNA (spacer regions) for all three species and chose a spacer length of approximately 20 bp to maximise target specificity. The Protospacer Adjacent Motif (PAM) region is identified by the sequence 5’-TTTV-3’ (V=A, G, C), and gRNA complimentary protospacer guanine (G)/cytosine (C) content between 30% and 60%. We used SnapGene (GSL Biotech; available at www.snapgene.com) to design the crRNAs for each species, which were synthesized by IDT. We used the same COI sequences in order to design the crRNA and RPA primers and confirmed the specificity of each crRNA using NCBI BLAST. For each species, two crRNAs were designed and tested (see *SI Appendix*, Table S2 for more details on crRNA design).

We also tested Cas12a nucleases, including *Lachnospiraceae bacterium ND2006* Cas12a (LbCas12a), *Acidaminococcus sp.* Cas12a (AsCas12a) and *Eubacterium rectale* Cas12a (ErCas12a), to determine trans-cleavage activity (*SI Appendix,* Table S4).

### Specificity assay

To demonstrate the specificity of our assay, we selected the RPA primer which amplified the COI region for three species and the Cas12a protein which showed the highest trans-cleavage activity. This optimised RPA-CRISPR-Cas12a assay was used with each gRNA to evaluate its specificity to purified PCR products of the target COI gene from *K. lycopersicella, P. absoluta*, and *S. atriplicella*. Specificity was assessed using the fluorescent-based assay as detailed below.

### Estimation of Limit of Detection (LoD)

Limit of detection for all three species was analysed with a series of eleven dilutions (10 nM, 1 nM, 100 pM, 10 pM, 1 pM, 100 fM, 10 fM, 1 fM, 100 aM, 10 aM and 1 aM) from PCR products. The PCR reaction was set up for species identification using COI primers, as described previously, and PCR products were cleaned up using the DNA Clean & Concentrator-5 kit (Zymo Research #D4014). We later quantified the concentration of PCR fragments using a Nanodrop (Thermo Scientific, ND-ONE-W) and a Qubit^TM^ Flex fluorometer (Thermo Fisher Scientific, Q33327) to prepare the above dilution series.

The LoD for the COI gene of each species was determined by amplifying the gene using RPA and then detecting it at the abovementioned eleven concentrations with our CRISPR assay. For LoD determination, the mean fluorescence was measured at 40 min and t=5 (*F*/*F* = 5) for 12 three replicates and compared to three control replicates that lacked target DNA. All analyses were done using GraphPad Prism (Version 9.4.1).

### Fluorescence-based detection assay

For the fluorescence-based detection assay, the Cas12a reaction was set up for a total volume of 40 μL (including activator/DNA) with 500 nM of Cas12a (LbCas12a, ErCas12a & AsCas12a) enzyme and 3 μM of crRNA in 10X NEBuffer 2.1. The solution was incubated at 37°C for 10 min before transferring to a 384-well plate containing 100 nM FAM fluorophore-quencher (FQ) reporter and 2 μL of activator (target DNA). Assays with a no-template control (NTC), which does not contain any template DNA, were also performed. Fluorescent excitation and emissions (λex: 485/20 nm, λem: 528/20 nm) resulting from Cas12a based trans-cleavage were measured every 2.5 min using a BioTek Synergy 2 microplate reader (Agilent). In a diagnostic specificity assay using field samples, fluorescence was also detected using the Analytik Jena UVP GelStudio PLUS system under blue light. For LoD experiments, a three-fold change in the ratio of fluorescence value of the sample to fluorescence value of the NTC is indicative of a positive result (48).

### Lateral flow detection assay

To evaluate the use of lateral flow readout, we screened different concentrations of the FAM-biotin reporter at 100, 200, and 300 nM and selected the highest concentration. For the lateral flow readouts, the CRISPR-Cas12a/RPA reaction was carried out by combining 500 nM of AsCas12a, 3 μM of crRNA, and 200 nM of FAM-Biotin reporter in 10X NEBuffer 2.1 for a volume of 48 μL, to which we later added 2 μL of the corresponding RPA product, which we incubated for 15 min at 37°C. A lateral flow strip (HybriDetect 1, Milenia Biotech) was then dipped into the reaction tube and a strip was checked after 5 min to determine the presence or absence of the target gene (*SI Appendix,* Table S8-9).

### Diagnostic specificity of CRISPR detection

To confirm whether our assay was effective for all three species, we conducted a blind test. A diagnostician received samples with codes and was unaware of sample details. Samples for this assay were prepared for the diagnostician from the pool of samples of *P. absoluta, K. lycopersicella* and *S. atriplicella* for each experiment. Three experiments were performed, with each consisting of 17 individuals of one of the target species, 6 randomly chosen members of the other two non-target species, and an NTC.

To detect all the insect samples, RPA was performed using genomic DNA extracts (*SI Appendix,* Table S6) followed by fluorescence and lateral flow reading, as explained previously.

### One-pot RPA-CRISPR-Cas12a detection

The RPA preamplification and CRISPR-Cas12a reaction were combined in one reaction. We adopted and modified the methods of Ding et al. (31), Sun et al. (32) and Xiong et al. (33). We prepared two separate components: an RPA mix, and a CRISPR-Cas12a mix. The RPA mix (25 µL) was prepared using the TwistAmp® Basic kit, which was similar to the RPA reaction in the two-pot system but with slight modifications. One RPA pellet from the TwistAmp® Basic kit was resuspended by mixing 29.5 µL of the rehydration buffer. After resuspension, 2.4 µL (10 µM) of each forward and reverse primer, 10 µL of nuclease-free PCR-grade water and 0.2 µL (100 nM) of Fluorophore-Quencher (FQ) was added, for a total final volume of 44.5 µL. After gentle vortexing, each 44.5 µL RPA mix was divided into two reactions of volume 22.25 µL and 1.5 μL of activator (target DNA) along with 1.25 µL (280 mM) of magnesium acetate are added making final volume to 25 µL. The CRISPR-Cas12a mix (5 µL) was prepared by mixing 1.25 µL 2000 nM AsCas12a enzyme, 0.85 µL 3000 nM of crRNA in 2.5 µL 10X NEBuffer 2.1, and 0.4 µL of nuclease-free PCR-grade water.

For the one-pot experiment, 25 µL of the RPA mix was gently placed at the bottom of a reaction tube and 5 µL of the CRISPR-Cas12a mix was added to the lid of the reaction tube before gently closing the lid. Then tube was incubated for 30 min at 37°C. The RPA and CRISPR-Cas12a mixes were combined with a short spin in a centrifuge. The tube containing the mixture was later placed in a CFX96 Real-Time PCR system with a C1000 Thermal Cycler module (Bio-rad) for fluorescence measurements for every 30 seconds per cycle for 60 cycles.

The sensitivity of the one-pot RPA-CRISPR-Cas12a detection assay was determined using a ten-fold serial dilution of template genomic DNA for all three species. The initial DNA concentration of 30 ng/μl was serially diluted from 10^−0^ ng/μl to 10^−10^ ng/μl and was used in the one-pot assay as described above. For diagnostic specificity of the one-pot detection assay, we used the same samples for a blind test, following the same protocol described in the diagnostic specificity assay of the two-pot CRISPR detection method.

### Statistical analysis

All data were processed and visualized using GraphPad Prism (Version 9.4.1) and figures were arranged in Adobe Illustrator. The number of replicates used for experiments are indicated in the figure legends. For fluorescence analyses of CRISPR reactions, the data were normalized by dividing all values by the initial value at *t*[=[5[min (*F*/*F*t =[5) for each replicate to allow for ∼5 min of temperature equilibration in the plate reader at the start of the assay and then calculated Standard error of the mean (SEM) for three replicates.

### Data Availability

All newly generated data in this study are included in the article and the *SI Appendix*.

## Acknowledgments

Authors acknowledge Dr. Tyler J. Wist, Research Scientist, Agriculture and Agri-Food Canada; Dr. Benjamin Lee, Postdoctoral researcher, University of California Davis, USA and Dr. A.S.R. Bajracharya, Scientist, Nepal Agricultural Research Institute, Nepal for sending specimens of *S. atriplicella*, *K. lycopersicella* and *P. absoluta*, respectively. Our sincere gratitude to Dr. Jean-Francois Landry, Research Scientist, Agriculture and Agri-Food Canada for providing adult images of *S. atriplicella* and *K. lycopersicella.* SRP expresses his sincere thanks to Dr. Ashok Kumar Singh, Director, Indian Agricultural Research Institute, New Delhi and Dr. Trilochan Mohapatra, Former Secretary, DARE and Director General, and Dr. D. K. Yadav, ADG (Seed), Indian Council of Agricultural Research, New Delhi, India for necessary permissions and their encouragement. We thank the National Biodiversity Authority, Government of India (File No. NBA/TechAppl./9/FormB/INBAB202203310/21/21-22/3984; dated:17.02.2022) for necessary approvals. PKJ was supported by NIH grant NIH-NIGMS R35GM147788. This research was supported by a cooperative agreement (#AP21PPQS&T00C030) between the USDA Animal and Plant Health Inspection Service and the University of Florida. Mention of trade names or commercial products in this publication is solely for the purpose of providing specific information and does not imply recommendation or endorsement by the U.S. Department of Agriculture, an equal opportunity employer.

## Data Availability

All the data supporting the findings of this study are available within the Article and Supplementary Files or can be obtained from the corresponding author, A.Y.K, upon reasonable request.

